# Contribution of single mutations to selected SARS-CoV-2 emerging variants Spike antigenicity

**DOI:** 10.1101/2021.08.04.455140

**Authors:** Shang Yu Gong, Debashree Chatterjee, Jonathan Richard, Jérémie Prévost, Alexandra Tauzin, Romain Gasser, Yuxia Bo, Dani Vézina, Guillaume Goyette, Gabrielle Gendron-Lepage, Halima Medjahed, Michel Roger, Marceline Côté, Andrés Finzi

**Affiliations:** Centre de Recherche du CHUM, QC H2X 0A9, Canada; Department of Microbiology and Immunology, McGill University, Montreal, QC H3A 0G4, Canada; Département de Microbiologie, Infectiologie et Immunologie, Université de Montréal, Montréal, QC H2X 0A9, Canada; Department of Biochemistry, Microbiology and Immunology, and Center for Infection, Immunity, and Inflammation, University of Ottawa, Ottawa, ON K1H 8M5, Canada; Laboratoire de Santé Publique du Québec, Institut Nationale de Santé Publique du Québec, Sainte-Anne-de-Bellevue, QC H9X 3R5, Canada

**Keywords:** Coronavirus, COVID-19, SARS-CoV-2, Spike glycoproteins, RBD, ACE2, temperature, variants of concern, vaccines

## Abstract

Towards the end of 2020, multiple variants of concern (VOCs) and variants of interest (VOIs) have arisen from the original SARS-CoV-2 Wuhan-Hu-1 strain. Mutations in the Spike protein are highly scrutinized for their impact on transmissibility, pathogenesis and vaccine efficacy. Here, we contribute to the growing body of literature on emerging variants by evaluating the impact of single mutations on the overall antigenicity of selected variants and their binding to the ACE2 receptor. We observe a differential contribution of single mutants to the global variants phenotype related to ACE2 interaction and antigenicity. Using biolayer interferometry, we observe that enhanced ACE2 interaction is mostly modulated by a decrease in off-rate. Finally, we made the interesting observation that the Spikes from tested emerging variants bind better to ACE2 at 37°C compared to the D614G variant. Whether improved ACE2 binding at higher temperature facilitates emerging variants transmission remain to be demonstrated.

## INTRODUCTION

Severe Acute Respiratory Syndrome Coronavirus 2 (SARS-CoV-2), the causative agent of COVID-19, remains an major public health concern, infecting over 185 million individuals and causing over 4 million deaths worldwide (Dong et al., 2020). The replication cycle of SARS-CoV-2 starts with viral attachment to the target cell and fusion between viral and cellular membranes. The viral entry process is mediated by the mature Spike (S) glycoprotein trimer which is composed of exterior S1 and transmembrane S2 subunits. The S1 subunit mediates attachment using its receptor-binding domain (RBD) to interact with the host protein angiotensin converting enzyme 2 (ACE2) (Hoffmann et al., 2020; Shang et al., 2020; Walls et al., 2019), while the S2 subunit governs the fusion between the viral and cellular membranes (Walls et al., 2020; Wrapp et al., 2020).The Spike is a major target of the cellular and humoral responses elicited by natural infection. Accordingly, the antigen used in currently approved vaccines is the stabilized form of the SARS-CoV-2 S glycoprotein. These vaccines use adenoviral vectors (Sadoff et al., 2021a; Voysey et al., 2021) or mRNA vaccine platforms to express S glycoprotein (Baden et al., 2020; Polack et al., 2020). The S glycoprotein was selected due to its high immunogenicity and safety profiles after extensive research (Jackson et al., 2020; Krammer, 2020; Mulligan et al., 2020; Sadoff et al., 2021b).

Although the approval of several vaccine platforms has given us hope to end the pandemic, the asymmetric distribution of doses between rich and poor countries and the rapid emergence of SARS-CoV-2 variants is preoccupying. The Spike is under high selective pressure to evade host immune response, improve ACE2 affinity, escape antibody recognition and achieve high transmissibility (Prévost and Finzi, 2021). The first identified D614G mutation in the Spike glycoprotein became dominant among the rapidly spreading emerging variants (Korber et al., 2020; Plante et al., 2021). In late 2020, several other variants emerged throughout the world, including the variants of concern (VOCs) B.1.1.7 (Alpha), B.1.351 (Beta), P.1 (Gamma) and B.1.617.2 (Delta), as well as the variants of interest (VOIs) B.1.429 (Epsilon), B.1.526 (Iota), B.1.617.1 (Kappa) and B.1.617 (CDC; Deng et al., 2021; ECDC, 2020, 2021; Ferreira et al., 2021; Mwenda M, 2021; West et al., 2021). Critical mutations providing a fitness increase became rapidly selected in most emerging variants. For example, the N501Y substitution that was first observed in the B.1.1.7 lineage and provides enhanced ACE2 binding (Prévost et al., 2021; Starr et al., 2020; Zhu et al., 2021), is now present in B.1.351, P.1, and P.3 lineages. Similarly, the E484K and K417N/T mutations in the RBD that were first described in the B.1.351 and P.1 lineages likely due to immune evasion from vaccine or natural infection-elicited antibodies (Amanat et al., 2021; Wang et al., 2021), are now present in several other lineages (Rambaut et al., 2020). Hence, it is important to closely monitor not only the emerging variants but also single mutations to better understand their contribution to replicative fitness and/or ability to evade natural or vaccine-induced immunity.

Here, by performing detailed binding and neutralization experiments with plasma from naturally infected and vaccinated individuals, we provide a comprehensive analysis of the antigenicity of the Spike from selected VOCs (B.1.1.7, B.1.351, P.1 and B.1.617.2) and VOIs (B.1.429, B.1.526, B.1.617, B.1.617.1).

## MATERIALS AND METHODS

### Ethics Statement

All work was conducted in accordance with the Declaration of Helsinki in terms of informed consent and approval by an appropriate institutional board. Blood samples were obtained from donors who consented to participate in this research project at CHUM (19.381). Plasmas were isolated by centrifugation.

### Plasmas and antibodies

Plasmas of SARS-CoV-2 naïve-vaccinated and previously infected pre- and post-first dose vaccination donors were collected, heat-inactivated for 1 hour at 56°C and stored at −80°C until use in subsequent experiments. Plasma from uninfected donors collected before the pandemic were used as negative controls in our flow cytometry and neutralization assays (not shown). The S2-specific monoclonal antibody CV3-25 was used as a positive control and to normalized Spike expression in our flow cytometry assays and was previously described (Jennewein et al., 2021; Mothes et al., 2021; Tauzin et al., 2021; Ullah et al., 2021). ACE2 binding was measured using the recombinant ACE2-Fc protein, which is composed of two ACE2 ectodomains linked to the Fc portion of the human IgG (Anand et al., 2020).Alexa Fluor-647-conjugated goat anti-human Abs (Invitrogen) were used as secondary antibodies to detect ACE2-Fc and plasma binding in flow cytometry experiments.

### Cell lines

293T human embryonic kidney cells (obtained from ATCC) were maintained at 37°C under 5% CO_2_ in Dulbecco’s modified Eagle’s medium (DMEM) (Wisent) containing 5% fetal bovine serum (VWR) and 100 µg/ml of penicillin-streptomycin (Wisent). The 293T-ACE2 cell line was previously described (Prévost et al., 2020) and was maintained in medium supplemented with 2 µg/mL of puromycin (Millipore Sigma).

### Plasmids

The plasmid encoding B.1.1.7, B.1.351, P.1, and B.1.526 Spikes were codon-optimized and synthesized by Genscript. Plasmids encoding B.1.617, B.1.617.1, B.1.617.2 Spikes were generated by overlapping PCR using a codon-optimized wild-type SARS-CoV-2 Spike gene (GeneArt, ThermoFisher) that was synthesized (Biobasic) and cloned in pCAGGS as a template. Plasmids encoding B.1.429, D614G and other SARS-CoV-2 Spike single mutations were generated using the QuickChange II XL site-directed mutagenesis protocol (Stratagene) and the pCG1-SARS-CoV-2-S plasmid kindly provided by Stefan Pöhlmann. The presence of the desired mutations was determined by automated DNA sequencing.

### Protein expression and purification

FreeStyle 293F cells (Invitrogen) were grown in FreeStyle 293F medium (Invitrogen) to a density of 1 x 10^6^ cells/mL at 37°C with 8 % CO_2_ with regular agitation (150 rpm). Cells were transfected with a plasmid coding for SARS-CoV-2 S RBD using ExpiFectamine 293 transfection reagent, as directed by the manufacturer (Invitrogen). One week later, cells were pelleted and discarded. Supernatants were filtered using a 0.22 µm filter (Thermo Fisher Scientific). The recombinant RBD proteins were purified by nickel affinity columns, as directed by the manufacturer (Invitrogen). The RBD preparations were dialyzed against phosphate-buffered saline (PBS) and stored in aliquots at −80°C until further use. To assess purity, recombinant proteins were loaded on SDS-PAGE gels and stained with Coomassie Blue.

### Virus neutralization assay

293T cells were transfected with the lentiviral vector pNL4.3 R-E-Luc (NIH AIDS Reagent Program) and a plasmid encoding for the indicated Spike glycoprotein (D614G, B.1.1.7, P.1, B.1.351, B.1.429, B.1.526, B.1.617, B.1.617.1, B.1.617.2) at a ratio of 10:1. Two days post-transfection, cell supernatants were harvested and stored at −80°C until use. 293T-ACE2 target cells were seeded at a density of 1×10^4^ cells/well in 96-well luminometer-compatible tissue culture plates (Perkin Elmer) 24h before infection. Pseudoviral particles were incubated with the indicated plasma dilutions (1/50; 1/250; 1/1250; 1/6250; 1/31250) for 1h at 37°C and were then added to the target cells followed by incubation for 48h at 37°C. Then, cells were lysed by the addition of 30 µL of passive lysis buffer (Promega) followed by one freeze-thaw cycle. An LB942 TriStar luminometer (Berthold Technologies) was used to measure the luciferase activity of each well after the addition of 100 µL of luciferin buffer (15mM MgSO_4_, 15mM KPO_4_ [pH 7.8], 1mM ATP, and 1mM dithiothreitol) and 50 µL of 1mM d-luciferin potassium salt (Prolume). The neutralization half-maximal inhibitory dilution (ID_50_) represents the plasma dilution to inhibit 50% of the infection of 293T-ACE2 cells by SARS-CoV-2 pseudoviruses.

### Cell surface staining and flow cytometry analysis

293T were transfected with full length SARS-CoV-2 Spikes and a green fluorescent protein (GFP) expressor (pIRES2-eGFP; Clontech) using the calcium-phosphate method. Two days post-transfection, 293T-Spike cells were stained with the CV3-25 Ab, ACE2-Fc or plasma from SARS-CoV-2-naïve or recovered donors. Briefly, 5 µg/mL CV3-25 or 20 µg/mL ACE2-Fc were incubated with cells at 37°C or 4 °C for 45 min. Plasma from SARS-CoV-2 naïve or convalescent donors were incubated with cells at 37°C. The percentage of Spike-expressing cells (GFP+ cells) was determined by gating the living cell population based on viability dye staining (Aqua Vivid, Invitrogen). Samples were acquired on a LSRII cytometer (BD Biosciences), and data analysis was performed using FlowJo v10.7.1 (Tree Star). The conformational-independent S2-targeting mAb CV3-25 was used to normalize Spike expression. CV3-25 was shown to be effective against all Spike variants (Mothes et al., 2021). The Median Fluorescence intensities (MFI) obtained with ACE2-Fc or plasma Abs were normalized to the MFI obtained with CV3-25 and presented as ratio of the CV3-25-normalized values obtained with the D614G Spike.

### Biolayer Interferometry

Binding kinetics were performed on an Octet RED96e system (FortéBio) at 25°C with shaking at 1,000 RPM. Amine Reactive Second-Generation (AR2G) biosensors were hydrated in water, then activated for 300 s with an S-NHS/EDC solution (Fortébio) prior to amine coupling. SARS-CoV-2 RBD proteins were loaded into AR2G biosensor at 12.5 µg/mL in 10mM acetate solution pH5 (Fortébio) for 600 s and then quenched into 1M ethanolamine solution pH8.5 (Fortébio) for 300 s. Baseline equilibration was collected for 120 s in 10X kinetics buffer. Association of sACE2 (in 10X kinetics buffer) to the different RBD proteins was carried out for 180 s at various concentrations in a two-fold dilution series from 500nM to 31.25nM prior to dissociation for 300 s. The data were baseline subtracted prior to fitting performed using a 1:1 binding model and the FortéBio data analysis software. Calculation of on-rates (K_a_), off-rates (K_dis_), and affinity constants (K_D_) was computed using a global fit applied to all data.

## RESULTS

### ACE2 recognition by SARS-CoV-2 single mutants and full Spike variants

Since the SARS-CoV-2 Spike is under strong selective pressure and is responsible for interacting with the ACE2 receptor, we measured the ability of the Spike from emerging variants to interact with ACE2 and the contribution of each single mutations toward this binding. Plasmids expressing the SARS-CoV-2 full Spike harboring single or combined mutations from emerging variants were transfected into HEK 293T cells. Spike expression was normalized with the conformationally independent, S2-specific CV3-25 monoclonal antibody (mAb) (Mothes et al., 2021; Ullah et al., 2021). ACE2 binding was measured using the recombinant ACE2-Fc protein, which is composed of two ACE2 ectodomains linked to the Fc portion of the human IgG (Anand et al., 2020). Alexa Fluor 647-conjugated secondary Ab was then used to detect ACE2-Fc binding to cell-surface Spike by flow cytometry. At the time of writing, B.1.617.1 and B.1.617.2 came into importance and only the full Spikes from these variants were synthesized, not the single mutations. When compared to the D614G Spike, all tested Spike variants, with the exception of B.1.617.1, presented significantly higher ACE2 binding (Fig 1A).

**Figure 1.**
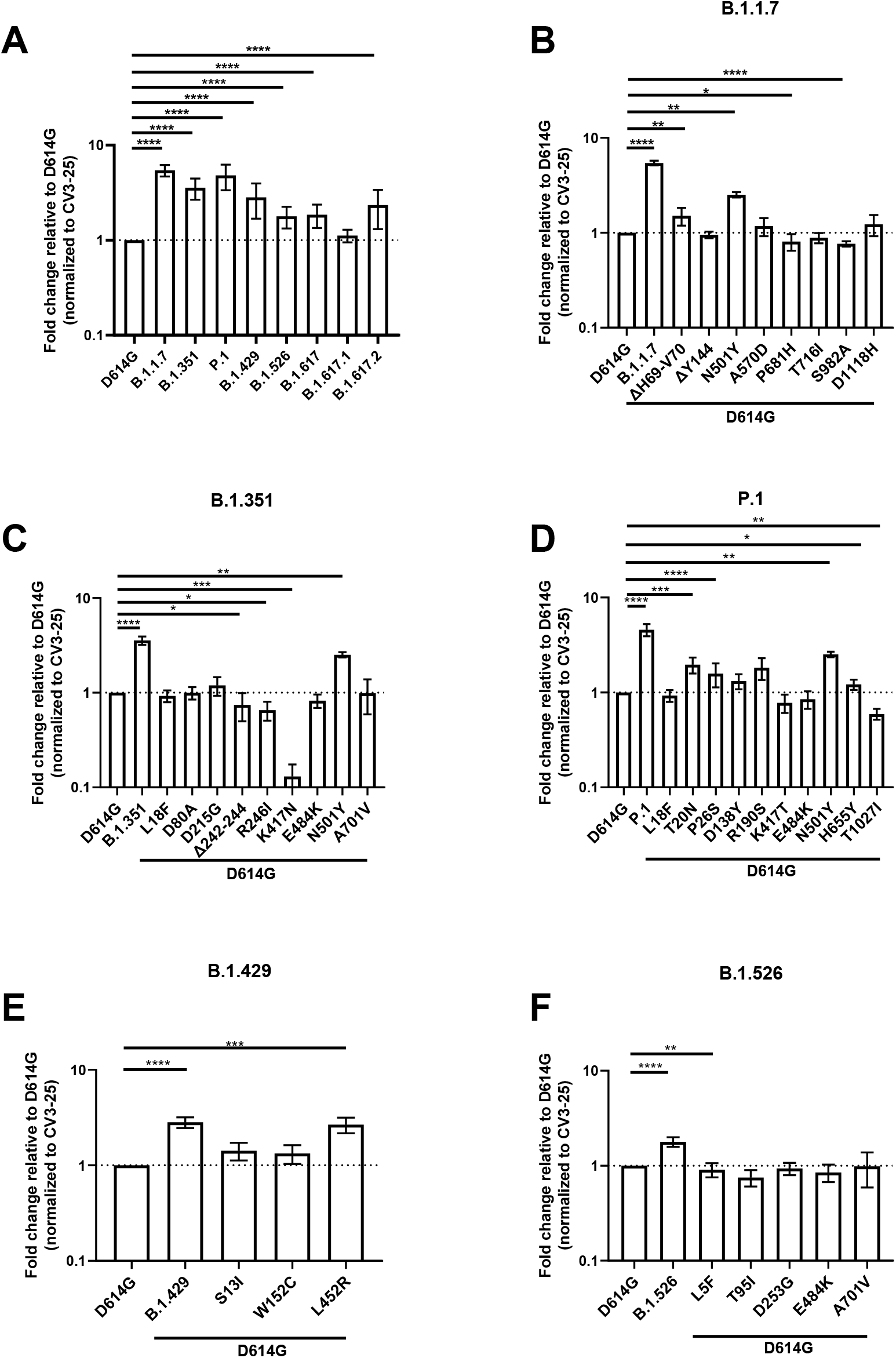
Evaluation hACE2 Fc binding to SARS-CoV-2 Spike variants. HEK 293T cells were transfected to express the indicated SARS-CoV-2 Spike variants. Two days post transfection, cells were stained with ACE2-Fc or with CV3-25 Ab as Spike expression control and analyzed by flow cytometry. ACE2-Fc binding to (A) full Spikes variants or the (B) B.1.1.7, (C) B.1.351, (D) P.1, (E) B.1.429, and (F) B.1.526 Spike and its corresponding single mutations are presented as a ratio of ACE2 binding to D614G Spike normalized to CV3-25 binding. Error bars indicate means ± SEM. Statistical significance has been performed using Mann-Whitney U test according to normality analysis (*p < 0.05; **p < 0.01; ***p < 0.001; ****p < 0.0001).

We then determine the contribution of individual Spike mutations on ACE2 binding to discern the ones contributing to the heightened receptor affinity of emerging variants. The B.1.1.7 Spike presented the highest ACE2-Fc interaction amongst all tested Spikes, which is a 5.43-fold increase in ACE2-Fc binding compared to D614G (Fig 1A-B and Table S2). The mutations that likely contribute to this phenotype are ΔH69-V70 in the N terminal domain (NTD) and N501Y in the RBD that enhanced binding by ∼1.51 and ∼2.52 folds, respectively (Fig 1B).

The Spike from B.1.351 also presented significantly higher ACE2-Fc binding compared to D614G. Similarly to B.1.1.7, the N501Y mutation likely plays an important role in this phenotype (Fig. 1C). Interestingly, three mutations/deletion in this VOC decreased the interaction with ACE2-Fc, namely R246I and Δ242-244 in the NTD, as well as K417N in the RBD. The NTD substitution R246I decreased ACE2-Fc binding by ∼1.52 folds, the Δ242-244 deletion by ∼1.35 folds, whereas K417N had a greater impact with a decreased binding of ∼7.7 folds relative to D614G (Fig 1C). Of note, the E484K mutation, also found in the RBD of other emerging variants (P.1 and B.1.526) did not significantly impact the ACE2-Fc interaction.

The Spike from P.1 presented a ∼4.24-fold increase in binding compared to D614G (Fig 1D). Few NTD mutations, namely T20N, P26S, D138Y, and R190S, likely contributed to the increase in ACE2 binding, with ∼2, ∼1.6, ∼1.3 and ∼1.8-fold increase compared to D614G, respectively. Like the above-mentioned VOCs, N501Y also likely played a role in enhanced ACE2-Fc interaction. Interestingly, the RBD mutation K417T and the S2 mutation T1027I decreased the ACE2-Fc by ∼1.3 and ∼1.7 folds respectively. The H655Y mutation, near the S1/S2 cleavage site, also slightly increased ACE2 interaction by ∼1.2 folds (Fig 1D).

The Spike from B.1.429 augmented ACE2-Fc interaction by ∼2.8 folds (Fig 1E). This VOI has two NTD mutations, S13I and W152C, both of which did not significantly impact this interaction. On the other hand, its RBD mutation, L452R, increased ACE2-Fc binding by ∼2.7 folds, suggesting its major contribution to the phenotype of this variant (Fig 1E).

Lastly, the Spike from B.1.526 showed a ∼1.8-fold increase over D614G (Fig 1F). None of the mutation appears to explain the phenotype observed with the full variant.

While our results identified some key mutations enhancing ACE2 interaction (i.e., N501Y,L452R and mutation/deletion in the NTD), the overall increased ACE2 affinity from any given variant appears to result from more than the sum of the effect of individual mutations composing this variant.

### Impact of selected mutations on the affinity of Spike RBD for ACE2

Next, we used biolayer interferometry (BLI) to measure the binding kinetics of selected RBD mutants to soluble ACE2 (sACE2). Biosensors were coated with recombinant RBD and put in contact with increasing concentration of sACE2 (Fig 2). In agreement with previous reports (Prévost et al., 2021; Zhu et al., 2021), the N501Y mutation present in B.1.1.7, B.1.351, and P.1 significantly decreased the off-rate (K_dis_) (from 6.88×10^−3^ to 1.49×10^−3^ 1/s), presenting a 3.88-fold increase in K_D_ compared to its wild-type counterpart (Fig 2A-B, Table S1). Substitution at position K417 (either N or T) present in B.1.351 or P.1 lineages accelerated the off-rate kinetics, resulting in a 0.75- and 0.66-fold decrease in binding affinity (Fig 2C-D, Table S1). Although both K417N and K417T presented a modest increase in the on-rate kinetic by ∼1.56 and ∼ 1.11 folds, the accelerated off-rate kinetics dictated the overall decrease affinity of these mutants. No major changes were observed for the E484K mutation (Fig 2F). Interestingly, the L452R mutant did not have a major impact in ACE2 affinity when tested in the context of recombinant monomeric RBD (Fig. 2G) but presented enhanced binding within the context of full-length membrane anchored Spike (Fig. 1E), indicating that the overall phenotype of a mutant (in this case enhanced binding) cannot always be recapitulated with the RBD alone. Altogether, our results show that at least for the RBD mutants tested here, ACE2 affinity is mostly dictated by the dissociation kinetics.

**Figure 2.**
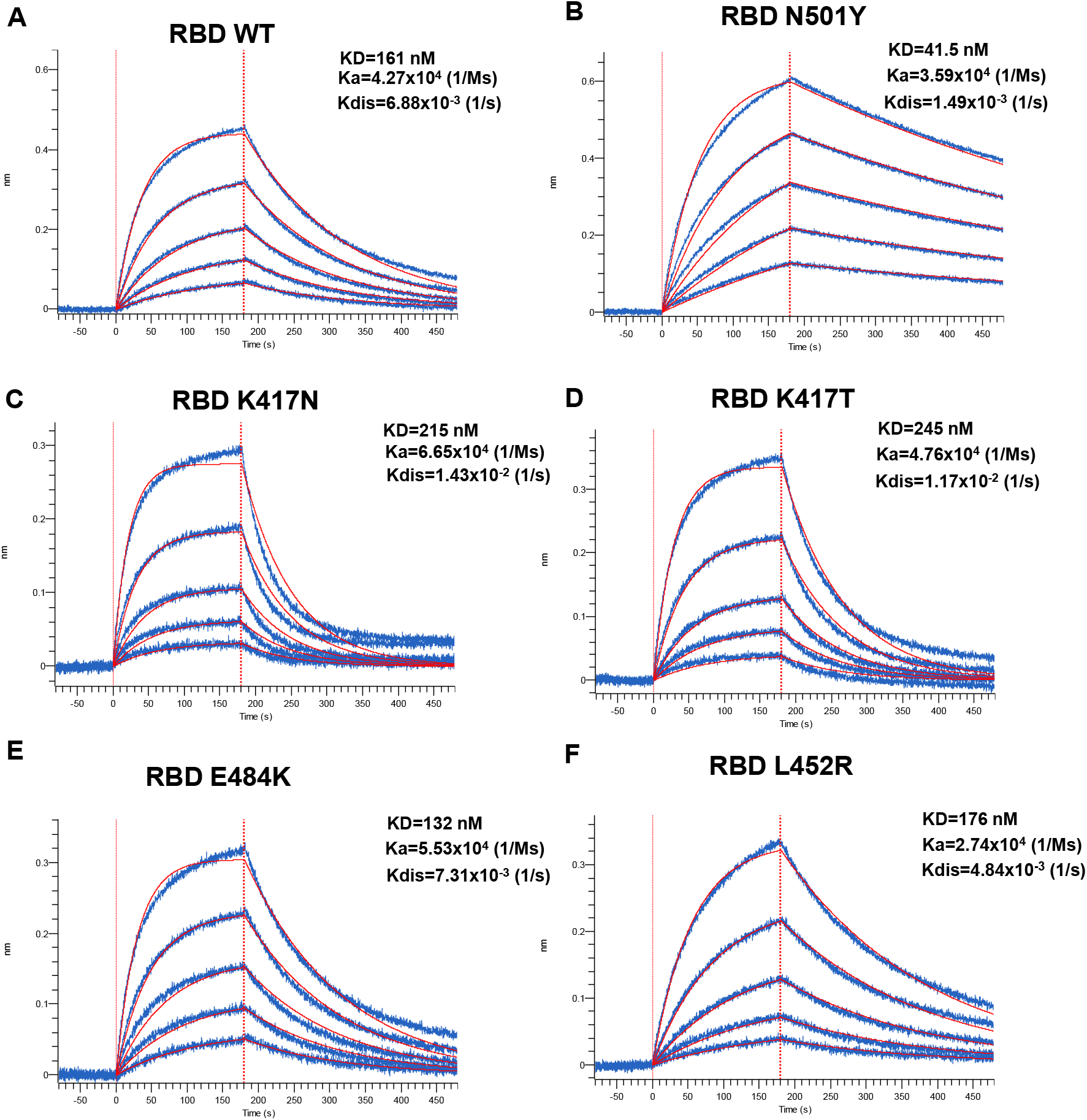
Kinetic Analysis of RBD interaction to hACE2 Binding by Biolayer Interferometry. Association of the different RBD proteins to sACE2 was carried out for 180s at various concentrations in a two-fold dilution series from 500nM to 31.25nM prior to dissociation for 300s for (A) WT, (B) N501Y, (C) K417N, (D) K417T, (F) E484K, and (G)L452R. Curve fitting was performed using a 1:1 binding model in the ForteBio data analysis software. Calculation of on-rates (K_a_), off-rates (K_dis_), and affinity constants (K_D_) was computed using a global fit applied to all data. Raw data are presented in blue and fitting models are in red. Results are summarized in Table S1.

### Effect of temperature on full variants Spike recognition of ACE2

It was recently shown that the affinity of Spike for ACE2 increases at low temperatures (Prévost et al., 2021). Interestingly, Prévost *et al*. also showed that the Spike from B.1.1.7 or harboring the N501Y mutation present better ACE2 binding at higher temperatures compared to the D614G strain (Prévost et al., 2021). To evaluate whether the Spikes from the emerging variants tested here also shared this phenotype, Spike-expressing cells were incubated at either 4°C or 37°C, and their ACE2-Fc binding was measured by flow cytometry. As presented in Fig. 3, the binding of ACE2-Fc to cell surface Spike was higher at cold temperature (4°C) compared to warm temperature (37°C) for all the variants. The impact of cold temperature on ACE2 binding was, however, more pronounced for the D614G Spike (3.62-fold increase) comparatively to Spikes from emerging variants (1.57 to 3.08-fold increase). While ACE2 displayed higher binding for the different emerging variants Spikes at 37°C, similar level of binding could only be achieved for the D614G Spike when decreasing the temperature to 4°C. This suggests that Spikes from emerging variants are able to bypass the temperature restraint to achieve high ACE2 binding. Interestingly, variants harboring the N501Y mutation (B.1.1.7, B.1.351 and P.1) exhibited an increase in ACE2 binding compared to D614G Spike at both 4°C and 37°C, while this phenotype was only observed at 37°C with Spike from the other variants. This indicate that the cold temperature and the N501Y mutation significantly impact Spike-ACE2 interaction.

**Figure 3.**
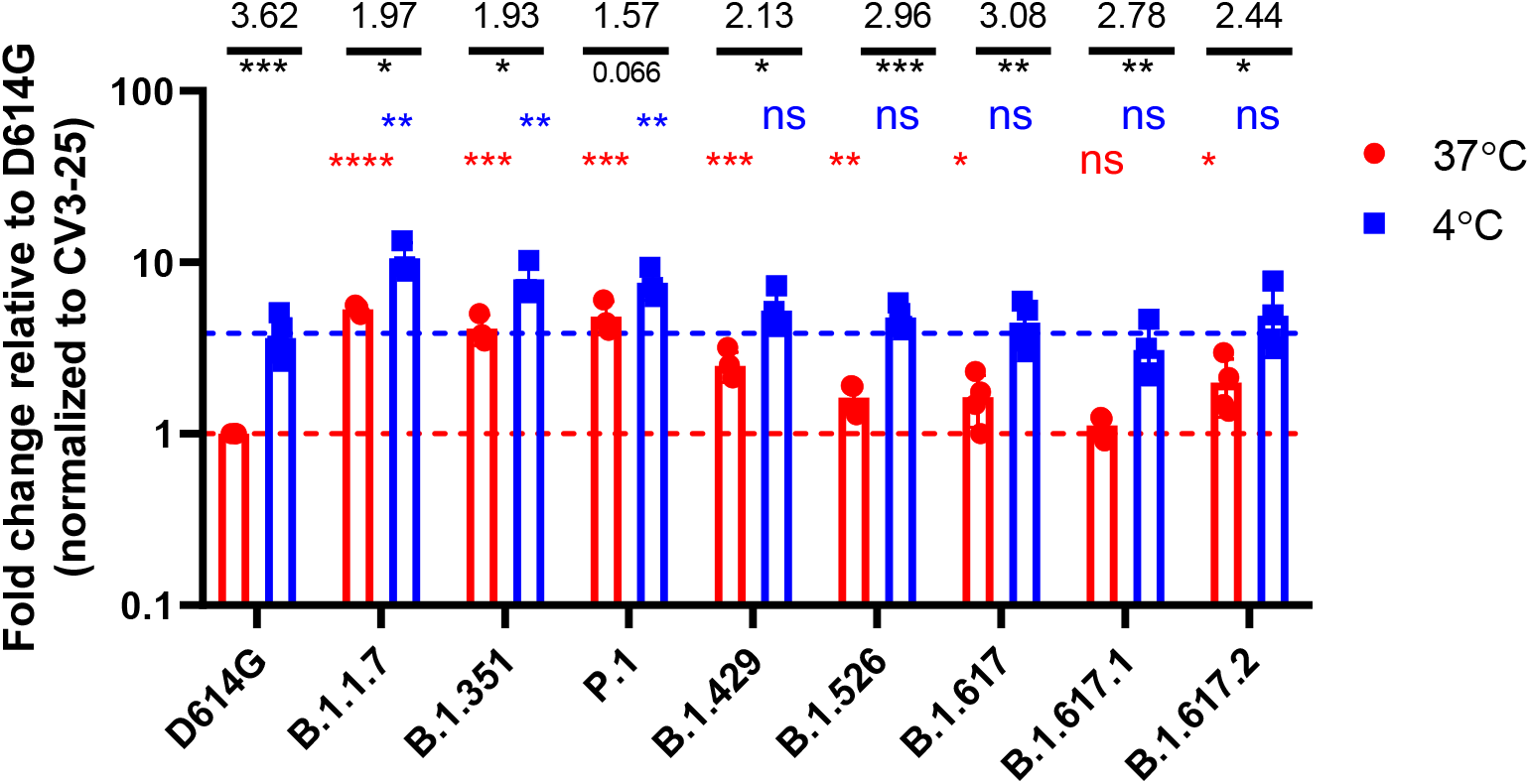
Evaluation of the impact of temperature on Spike-ACE2 interaction. HEK 293T cells were transfected with the indicated SARS-CoV-2 Spike variants. Two days post transfection, cells were stained with ACE2-Fc or with CV3-25 Ab as Spike expression control at 4°C or 37°C and analyzed by flow cytometry. ACE2-Fc binding to the different Spike variants are presented as a ratio of ACE2 binding to D614G Spik, normalized to CV3-25 binding at 37°C (red) or at 4°C (blue). Statistical analyses were used to compare each Spike at 4°C vs 37°C (black) or to compare variants Spike to D614G at 37°C (red) or at 4°C (blue). Fold changes of ACE2 binding at 4°C vs 37°C for each Spike is shown in black. Error bars indicate means ± SEM. Statistical significance has been performed using Mann-Whitney U test (*p < 0.05; **p < 0.01; ***p < 0.001; ****p < 0.0001, ns; non-significant).

### Recognition of Spike variants by plasma from vaccinated individuals

To gain information related to the antigenic profile of each emerging variants Spike and their single mutants, we used plasma collected three weeks post BNT162b2 vaccination from SARS-CoV-2 naïve (Fig. 4) or previously-infected individuals (Fig. 5) (Tauzin et al., 2021). HEK 293T cells were transfected with Spike from emerging variants or their individual mutants and plasma binding was evaluated by flow cytometry, as previously reported (Anand et al., 2021; Beaudoin-Bussières et al., 2020; Gasser et al., 2021; Prévost et al., 2020; Tauzin et al., 2021). When compared to D614G, Spike from B.1.1.7, P.1 and the recently emerged B.1.617.1 and B.1.617.2 variants were significantly less recognized by the plasma from SARS-CoV-2 naïve individuals (Fig 4A).

**Figure 4.**
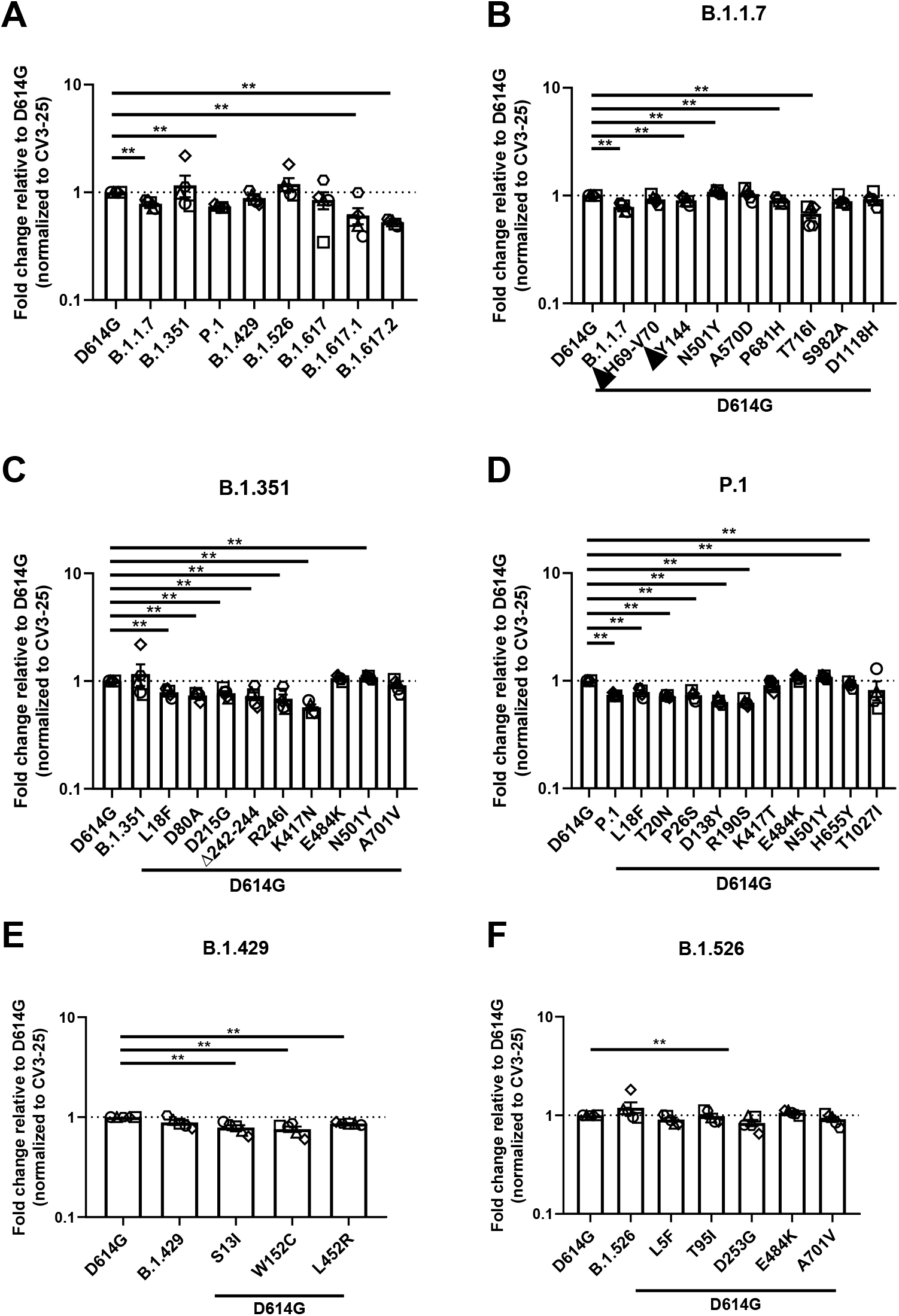
Recognition of SARS-CoV-2 Spike variants and single mutants by plasma from vaccinated SARS-CoV-2 naïve individuals. HEK 293T cells were transfected with the indicated SARS-CoV-2 spike variants. Two days post transfection, cells were stained with 1:250 dilution of plasma collected from naive post vaccinated individuals (n=3-5) or with CV3-25 Ab as control and analyzed by flow cytometry. Plasma recognition of (A) full Spike variants (B) B.1.1.7, (C) B.1.351, (D) P.1, (E) B.1.429, (F) B.1.526 Spike and variant-specific Spike single mutations are presented as ratio of plasma binding to D614G Spike normalized CV3-25 binding. Error bars indicate means± SEM. Statistical significance has been performed using Mann-Whitney U test (*p < 0.05; **p < 0.01; ***p < 0.001; ****p < 0.0001).

**Figure 5.**
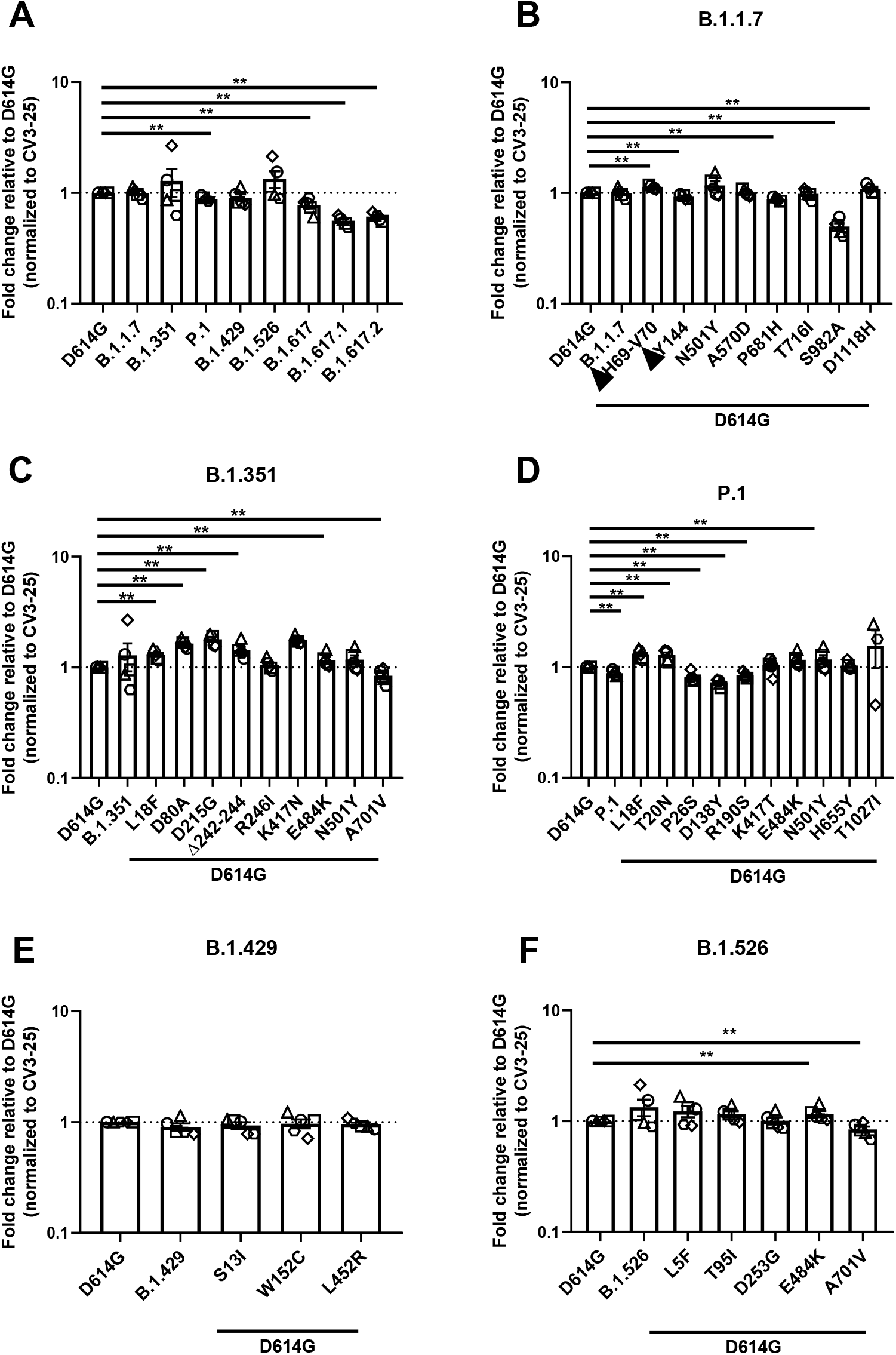
Recognition of SARS-CoV-2 Spike variants and single mutants by plasma from vaccinated previously-infected individuals. HEK293T cells were transfected with SARS-CoV-2 full Spike variants and stained with plasma collected 3 weeks post-first dose vaccinated previously infected individuals (n=3-5) or with CV3-25 Ab and analyzed by flow cytometry. Plasma recognition of (A) full Spike variants or the (B) B.1.1.7, (C) B.1.351, (D) P.1, (E) B.1.429, (F) B.1.526 Spikes and Spikes with their corresponding single mutations are presented as a ratio of plasma binding to D614G Spike normalized with CV3-25 binding. Error bars indicate means ± SEM. Statistical significance has been performed using Mann-Whitney U test (*p < 0.05; **p < 0.01; ***p < 0.001; ****p < 0.0001).

Mutations apparently contributing to the reduction of plasma recognition of the B.1.1.7 Spike are the ΔY144 deletion in the NTD, P681H and T716I near the S1/S2 cleavage site (Fig 4B). The N501Y substitution, also present in other emerging variants (B1.351 and P.1), is the only mutation that increased plasma recognition.

While we did not observe a significant decrease in plasma recognition of the B.1.351 Spike, most single mutants of this VOC (L18F, D80A, D215G, Δ242-244, R246I in the NTD, and K417N in the RBD) exhibit decreased binding. Of note, the K417N mutation that reduced ACE2 binding (Fig 1C), also significantly impacted plasma recognition by ∼1.75-fold compared to D614G (Fig 4C).

Polyclonal recognition of the P.1 Spike was reduced by ∼1.33 folds (Fig 4D). Our results show that all the NTD mutations, namely L18F, T20N, P26S, D138Y and R190S, attenuated the binding of naïve-vaccinated plasma Abs (Fig 4D). Furthermore, H655Y also contributed to the immuno-evasive phenotype of the full Spike (Fig 4D). Again, N501Y is the only mutation that increased plasma recognition, indicating its major role amongst all the mutations of this variant.

Although the full B.1.429 Spike did not show a significant evasion of plasma recognition, all of its mutations presented a significant decrease in plasma binding (Fig 4E). Both its NTD mutations, S13I and W152C, were less efficiently recognized by plasma compared to D614G (Fig 4E). The RBD mutation L452R also presented a minor ∼1.16-fold decrease in recognition by plasma from vaccinated individuals. Lastly, the B.1.526 full Spike variant did not significantly affect vaccine-elicited plasma recognition. The same phenotype is applicable to most of its mutations, with the exception of D253G substitution in the NTD, which showed a modest ∼1.2-fold decrease in binding (Fig 4F).

We then evaluated the recognition of our panel of Spikes (VOC, VOI and single mutants) by plasma from previously-infected vaccinated individuals (plasma recovered three weeks post vaccination), as recently described (Tauzin et al., 2021). When comparing all Spikes from emerging variants, the convalescent plasma pre- and post-first dose vaccination both effectively recognized all tested Spikes (Fig 5A, S1 and S2). Vaccinated convalescent individuals developed Abs that were able to robustly recognize and bind to the emerging variants B.1.1.7, B.1.351, B.1.429 and B.1.526 at a similar level than D614G. Binding was decreased by ∼ 1.14 folds for P.1, ∼1.3 folds for B.1.617 and by ∼ 1.8 and ∼1.64 folds for B.1.617.1 and B.1.617.2 Spikes, respectively (Fig 5A).

Examining each variant and their single mutations more closely, though full B.1.1.7 Spike did not significantly reduce plasma binding in convalescent post-vaccinated individuals, three of its single mutations, ΔY144, P681H, and S982A significantly affected plasma recognition. Substitution S982A in the S2 showed the most important reduction, by ∼2 folds compared to D614G (Fig 5B). Inversely, the deletion ΔH69-V70 and the substitution D1118H slightly enhanced the recognition of this Spike by plasma from previously-infected vaccinated individuals. However, combined together, the B.1.1.7 was recognized similarly to its D614G counterpart by these plasmas.

The B.1.351 Spike was efficiently recognized by plasma from previously-infected vaccinated individuals with a single mutation presenting lower detection (A701V) (Fig 5C). In the P.1 Spike we observed mutations in the NTD that decreased recognition (P26S, D138Y, and R190S) (Fig 5D). The Spikes from B.1.429 and B.1.526 were also efficiently recognized. Only the A701V substitution present in the B.1.526 reduced plasma binding (Fig.5F). Among all tested emerging variants, the Spikes from B.1.617.1 and B.1.617.2 presented the most important decrease in recognition by ∼1.8 and ∼1.64 folds compared to D614G (Figure 5A)

Altogether, our results highlight the difficulty in predicting the phenotype of a particular variant based on the phenotype of its individual mutations.

### Neutralization of Spike variants by plasma from vaccinated individuals

We then determined the neutralization profile of the different emerging variants Spikes using a pseudoviral neutralization assay (Anand et al., 2021; Beaudoin-Bussières et al., 2020; Ding et al., 2020; Gasser et al., 2021; Prévost et al., 2020). For this assay we used plasma from five individuals previously infected with SARS-CoV-2 (on average 8 months post-symptom onset) after one dose of the Pfizer/BioNtech BNT162b2 vaccine (samples were collected 22 days after immunization) (Tauzin et al., 2021). As we described previously (Tauzin et al., 2021), we incubated serial dilutions of plasma with pseudoviruses bearing the different Spikes before adding to HEK 293T target cells stably expressing the human ACE2 receptor (Tauzin et al., 2021). We obtained robust neutralization for all pseudoviral particles, with a modest, but significant decrease in neutralization against pseudoviruses bearing the Spike from B.1.351, B.1.526 and B.1.617.2 lineages, indicating that plasma from previously-infected individuals following a single dose have a relatively good neutralizing activity against emerging variants (Fig 6).

**Figure 6.**
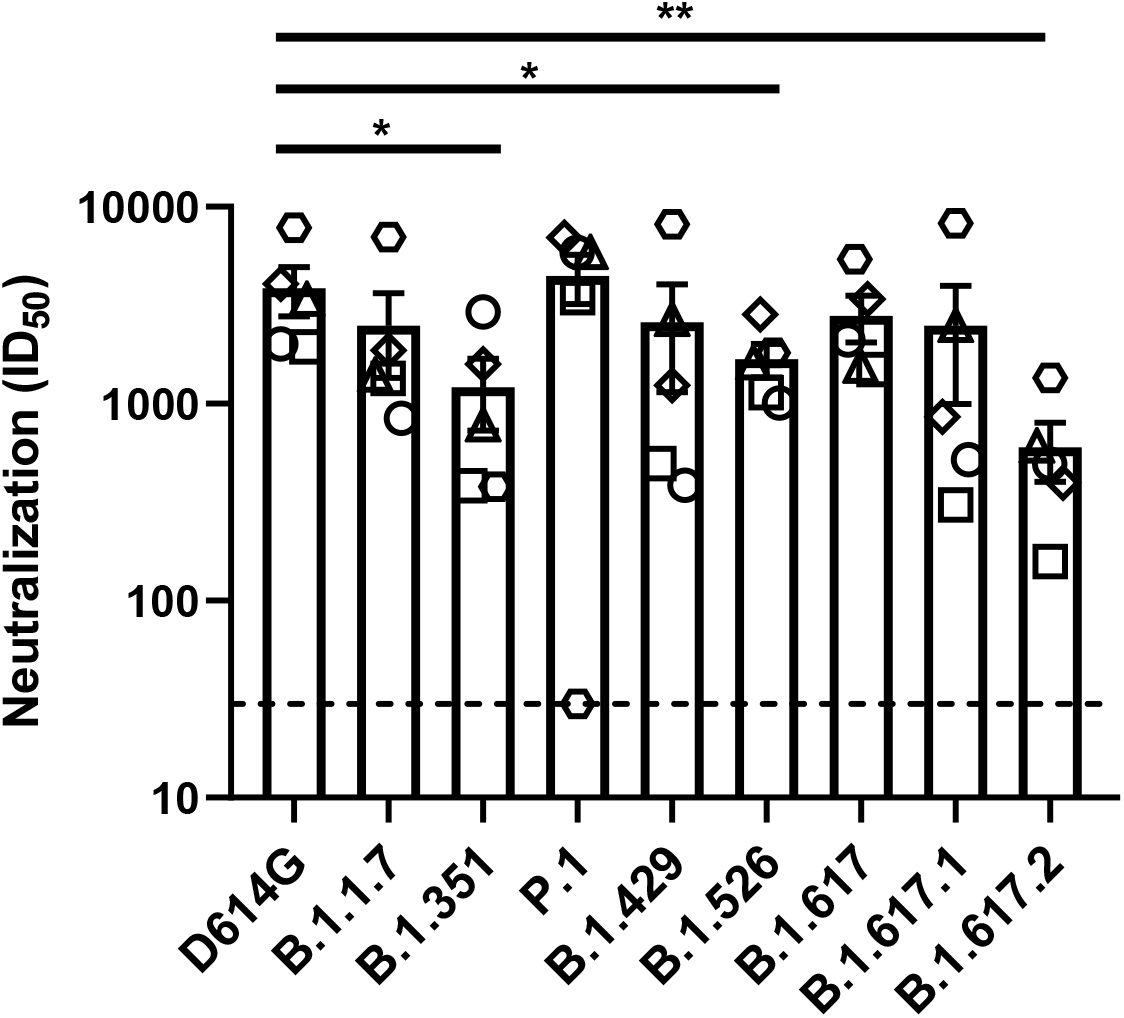
Neutralization of SARS-CoV-2 Spike variants by plasma from previously infected vaccinated individuals. Neutralizing activity of previously infected vaccinated individuals against pseudoviruses bearing the SARS-CoV-2 Spike variants were assessed. Pseudoviruses with serial dilutions of plasma were incubate for 1 h at 37°C before infecting 293T-ACE2 cells. ID_50_ against pseudoviruses were calculated by a normalized non-linear regression using GraphPad Prism software. Detection limit is indicated in the graph (ID_50_=50). Statistical significance has been performed using Mann-Whitney U test (*p < 0.05).

## DISCUSSION

In our study, we offer a comparative view examining the ACE2 binding properties of selected circulating variants, and the impact of their single mutations on plasma binding. We observed that Spikes from B.1.1.7, B.1.351, P.1, B.1.429, B.1.526, B.1.617, and B.1.617.2 lineages present increased ACE2 interaction. Consistent with previous reports (Leung et al., 2021; Starr et al., 2020; Washington et al., 2021), the N501Y mutation shared by B.1.1.7, B.1.351, and P.1 variants presented a significant increase for ACE2-Fc binding. While N501Y plays a major role in enhanced transmissibility and infectivity (Liu et al., 2021), variants which do not share this mutation have also gained the increased ACE2 binding by harboring other mutations, such as in the B.1.429 lineage, where the L452R showed higher ACE2 binding.

We also analyzed the impact of temperature in modulating the capacity of Spikes from emerging variants to interact with the viral receptor ACE2. For almost all tested Spikes, we observed a significant increase in ACE2 binding at cold temperature (4 °C). As recently reported, this could be explained by favorable thermodynamics changes allowing the stabilization of the RBD-ACE2 interface and by modulating the Spike trimer conformation (Prévost et al., 2021). While the D614G Spike necessitates lower temperature for optimal ACE2 interaction, Spikes from the different VOCs and VOIs seem to bypass this requirement to efficiently interact with ACE2 at higher temperature (37 °C). However, whether improved ACE2 binding at higher temperature facilitates emerging variants transmission and propagation remain to be demonstrated. Interestingly, variants harboring the N501Y mutation displayed improve ACE2 interaction compared to the D614G Spike, independently of the temperature, highlighting the critical impact of this substitution in improving Spike – ACE2 interaction. This reveals the importance of closely monitoring the appearance of this mutation among the current and future emerging variants. The appearance of this substitution could potentially impact the transmission and propagation of recent rapidly spreading emerging variants (such as the B.1.617.2 lineage) that do not harbor this mutation, by enhancing the affinity of their Spike for the ACE2 receptor at cold and warm temperatures.

We also found that plasma from vaccinated SARS-CoV-2 naïve and prior-infected individuals efficiently recognized the Spikes from emerging variants. However, as previously shown, plasma from vaccinated previously-infected individuals presented a higher and more robust recognition of all tested Spikes (Fig. S2)(Lucas et al., 2021; Tauzin et al., 2021). Accordingly, plasma from these individuals were able to neutralize pseudoviral particles bearing the different emerging variants Spikes, further highlighting the resilience of the deployed vaccines, which were based on the original Wuhan strain.

## CONCLUSIONS

Altogether, our results highlight the difficulty in predicting the phenotype of an emerging variant’s Spike, either related to ACE2 interaction, antigenic profile, infectivity and transmission based on the sum of the phenotype of single mutants making that particular Spike. Antigenic drift has been and remains a concern of the current pandemic (Callaway, 2021; Prévost and Finzi, 2021) and therefore, closely monitoring the functional properties of emerging variants remains of the utmost importance for vaccine design and to inform public health authorities to better manage the epidemic by implementing preventive interventions to control the spread of highly transmissible virus, and tailoring vaccination campaign.

### CRediT authorship contribution statement

**Shang Yu Gong:** Conceptualization, Methodology, Validation, Formal analysis, Investigation, Resources, Writing – original draft, Visualization. **Debashree Chatterjee:** Conceptualization, Methodology, Validation, Formal analysis, Investigation, Resources, Writing – original draft, Visualization. **Jonathan Richard:** Conceptualization, Methodology, Validation, Formal analysis, Investigation, Writing – original draft, Supervision. **Jérémie Prévost:** Conceptualization, Methodology, Resources. **Alexandra Tauzin:** Methodology, Resources, Validation, Formal analysis, Investigation. **Romain Gasser**: Methodology, Resources, Validation, Formal analysis, Investigation. **Dani Vézina:** Resources. **Guillaume Goyette:** Resources. **Gabrielle Gendron-Lepage**: Resources. **Michel Roger:** Resources, Writing - Review & Editing. **Marceline Côté:** Methodology, Writing - Review & Editing, Supervision, Funding acquisition. **Andrés Finzi**: Conceptualization, Methodology, Writing – original draft, Writing – original draft, Visualization, Project administration, Supervision, Funding acquisition.

## Acknowledgements

The authors thank the CRCHUM BSL 3 and Flow Cytometry Platforms for their technical assistance and Dr Sandrine Moreira (Laboratoire de Santé Publique du Québec) for helpful discussions. We thank Dr. Stefan Pöhlmann and Dr. Markus Hoffmann (Georg-August University) for the plasmids coding for SARS-CoV-2 WT Spike (Wuhan-Hu-1 strain) and Dr Jason McLellan for the plasmid coding for sACE2 expressor. The CV3-25 antibody was produced using the pTT vector kindly provided by the Canada Research Council.

## SUPPLEMENTAL TABLE

**Table S1.** Binding Kinetics of the interaction between SARS-CoV-2 RBD and sACE2 quantified by Biolayer Interferometry.

| RBD | ACE2 binding<br>(RBD/biolayer interferometry) |  |  |  |  |  |
| --- | --- | --- | --- | --- | --- | --- |
|  | KD (nM) | Fold change | Ka (1/Ms) | Fold change | Kdis (1/s) | Fold change |
| WT | 161 | 1 | 4.27x10<sup>4</sup> | 1 | 6.88x10<sup>-3</sup> | 1 |
| N501Y | 41.5 | 3.88 | 3.59x10<sup>4</sup> | 0.84 | 1.49x10<sup>-3</sup> | 0.22 |
| K417N | 215 | 0.75 | 6.65x10<sup>4</sup> | 1.56 | 1.43x10<sup>-2</sup> | 2.07 |
| K417T | 245 | 0.66 | 4.76x10<sup>4</sup> | 1.11 | 1.17x10<sup>-2</sup> | 1.70 |
| E484K | 132 | 1.22 | 5.53x10<sup>4</sup> | 1.29 | 7.31x10<sup>-3</sup> | 1.06 |
| L452R | 176 | 0.91 | 2.74x10<sup>4</sup> | 0.64 | 4.84x10<sup>-3</sup> | 0.70 |

**Table S2.**
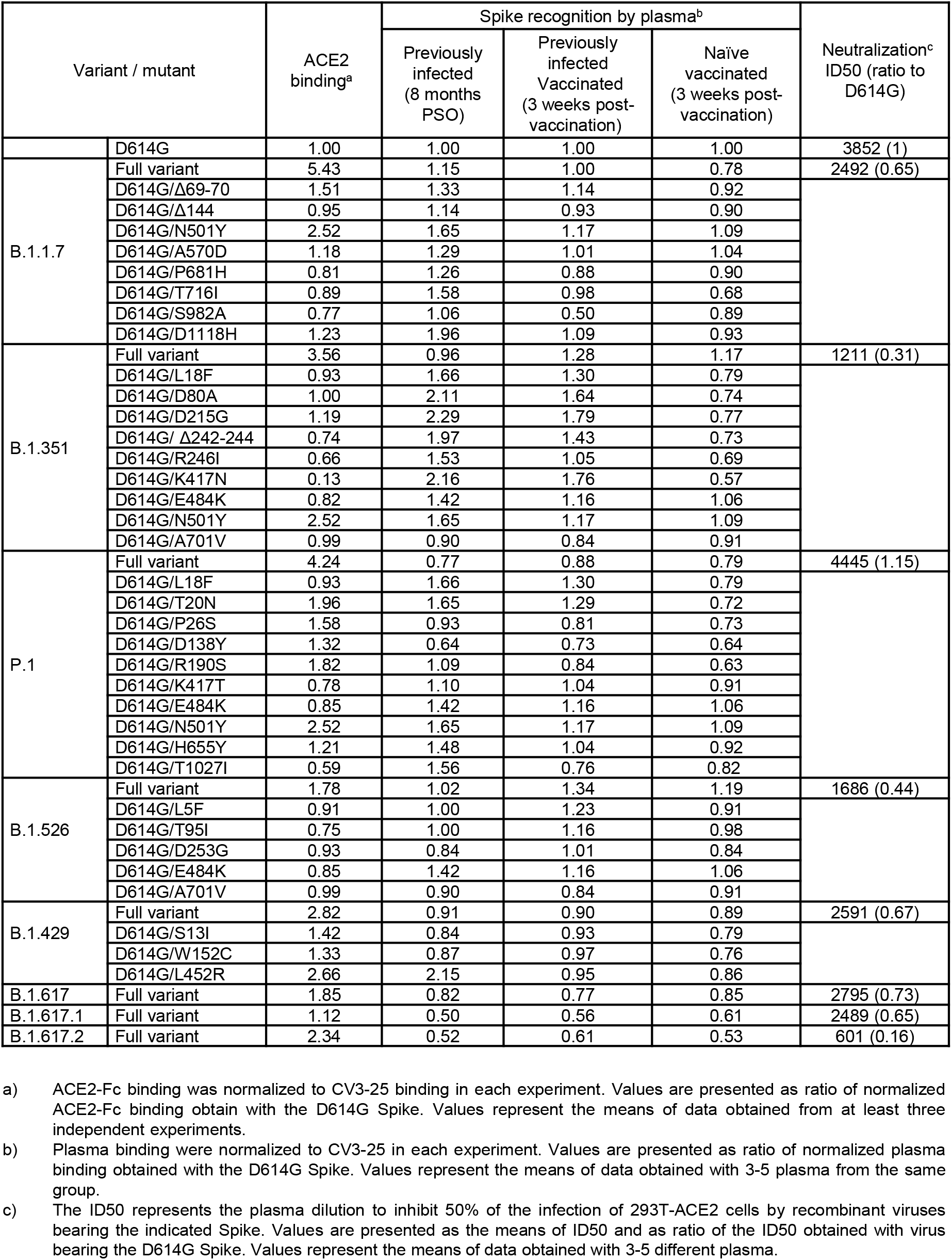
Summary of ACE2 Binding, Plasma binding, and Neutralization to Spike variants.

## SUPPLEMENTAL FIGURES

**Figure S1.**
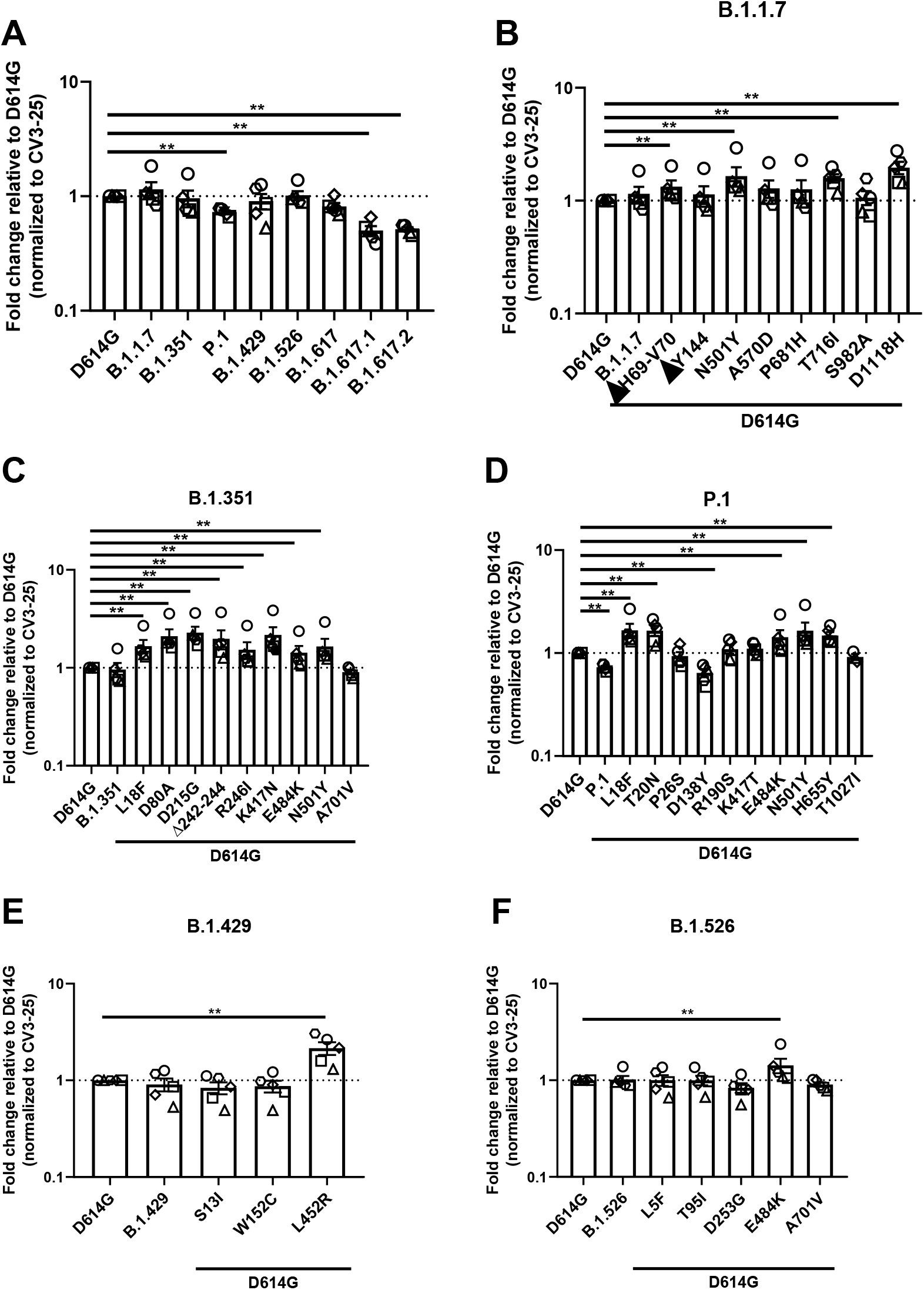
Recognition of SARS-CoV-2 Spike variants and single mutants by plasma from convalescent donors. HEK293T cells were transfected with the indicated SARS-CoV-2 spike variants and stained with plasma collected from individuals that were infected around 9 months before plasma collection (n=5) or with CV3-25Ab and analyzed by flow cytometry. Plasma recognition of (A) full Spike variants or the (B) B.1.1.7, (C) B.1.351, (D) P.1, (E) B.1.429, (F) B.1.526 Spikes and lineage specific Spike single mutations are presented as ratio of plasma binding to D614G normalized with CV3-25 binding. Error bars indicate means± SEM. Statistical significance has been performed using Mann-Whitney U test (*p < 0.05; **p < 0.01; ***p < 0.001; ****p < 0.0001).

**Figure S2.**
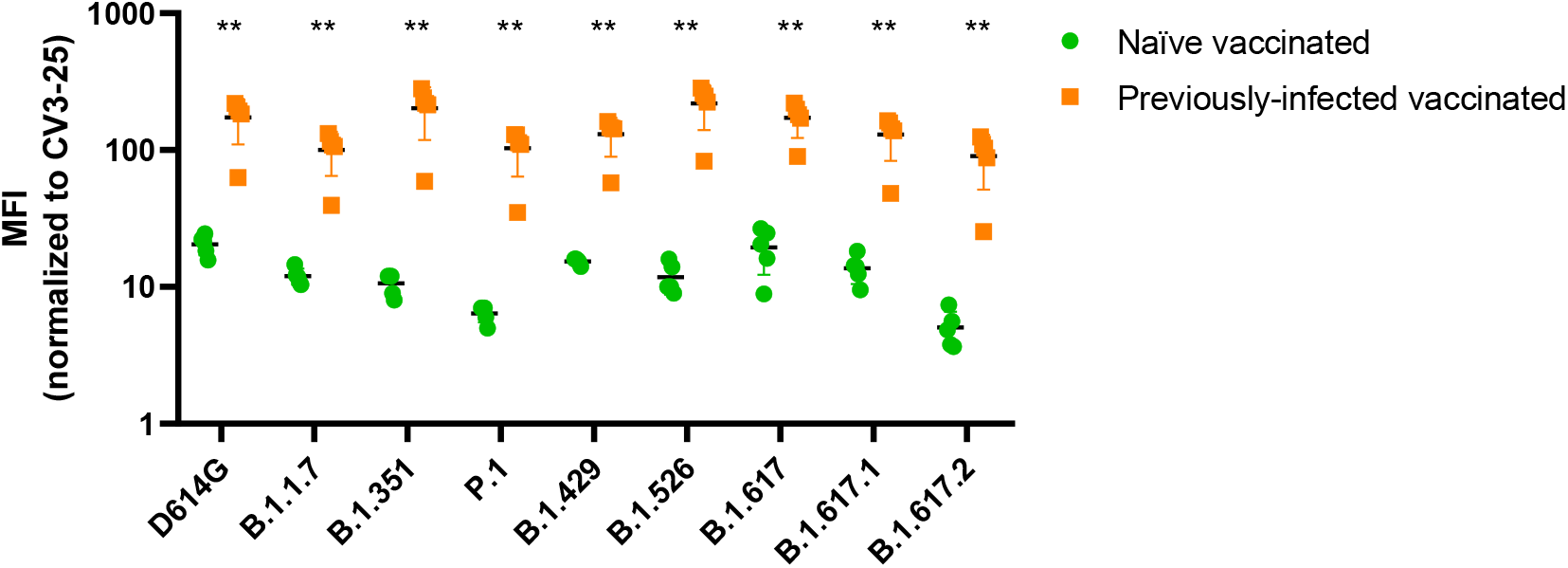
Recognition of SARS-CoV-2 Spike variants by plasma from naïve or previously-infected, vaccinated individuals. HEK293T cells were transfected with SARS-CoV-2 full Spike variants and stained with plasma collected 3 weeks post-first dose vaccinated previously infected individuals (n=5), first dose vaccinated naïve individuals (n=5), or CV3-25 Ab and analyzed by flow cytometry. Plasma recognition of full Spike variants are represented as Median Fluorescent Intensities (MFIs) normalized as a percentage of CV3-25 binding. Error bars indicate means ± SEM. Statistical significance has been performed using Mann-Whitney U test (*p < 0.05; **p < 0.01; ***p < 0.001; ****p < 0.0001).

